# Membrane-dependent actin polymerization mediated by the *Legionella pneumophila* effector protein MavH

**DOI:** 10.1101/2023.01.24.525393

**Authors:** Qing Zhang, Min Wan, Yuxin Mao

## Abstract

*L. pneumophila* propagates in eukaryotic cells within a specialized niche, the *Legionella*-containing vacuole (LCV). The infection process is controlled by over 330 effector proteins delivered through the type IV secretion system. In this study, we report that the *Legionella* MavH effector harbors a lipid-binding domain that specifically recognizes PI(3)P (phosphatidylinositol 3-phosphate) and localizes to endosomes when ectopically expressed. We show that MavH recruits host actin capping proteins (CP) and actin to the endosome via its CP interacting (CPI) motif and WH2-like actin-binding domain, respectively. In vitro assays revealed that MavH stimulates robust actin polymerization only in the presence of PI(3)P-containing liposomes and the recruitment of CP by MavH negatively regulates F-actin density at the membrane. Furthermore, in *L. pneumophila*-infected cells, MavH can be detected around the LCV at the very early stage of infection. Together, our results reveal a novel mechanism of membrane-dependent actin polymerization catalyzed by MavH that may play a role at the early stage of *L. pneumophila* infection by regulating host actin dynamics.

## INTRODUCTION

The gram-negative bacterium *Legionella pneumophila* is a facultative intracellular pathogen. Human infection, which occurs when aerosols contaminated by *Legionella* are inhaled, is found to be responsible for a severe form of pneumonia in humans known as Legionnaires’ disease (Fraser et al., 1977; McDade et al., 1977). *L. pneumophila* secretes over 330 effector proteins into host cells via its Dot/Icm (Defective Organelle Trafficking/Intracellular Multiplication) apparatus during infection (Burstein et al., 2009; Huang et al., 2011; Zhu et al., 2011). These proteins modulate every step in the infection process, including host cell entry (Hilbi et al., 2001; Watarai et al., 2001), maturation of a replication-competent *Legionella* containing vacuole (LCV) (Mondino et al., 2020), evading phagolysosomal fusion (Roy et al., 1998) and autophagy (Choy et al., 2012; Omotade and Roy, 2020), and final egress from the host cell (Flieger et al., 2018). Although the biological functions of many effectors have been elucidated, the exact molecular mechanisms of most effectors remain uncharacterized.

Actin is one of the most conserved proteins throughout evolution and exists in two distinct forms, the monomeric G-actin form, and the filamentous F-actin form. F-actin is highly dynamic with a net association of ATP-actin to the barbed (+) end and dissociation of ADP-actin monomers from the pointed (−) end (Pollard, 2016). The rapid assembly and disassembly of the actin cytoskeleton play an essential role in diverse cellular processes (Dominguez and Holmes, 2011), including phagocytosis, micropinocytosis, endocytosis, vesicle trafficking, cell motility, polarity, and cytokinesis. The dynamics of the actin cytoskeleton are tightly regulated by a large number of actin-binding proteins, such as actin nucleators, capping proteins, severing proteins, etc (Pollard, 2016; Pollard and Borisy, 2003). *de novo* F-actin assembly requires actin nucleators to overcome kinetic energy barriers (Rottner et al., 2017). Three major classes of nucleators have been identified so far: the Arp2/3 complex (Goley and Welch, 2006); the formins (Breitsprecher and Goode, 2013); and the tandem actin-binding domain proteins, such as Spire (Kerkhoff, 2006), Cobl (Ahuja et al., 2007), which promote actin nucleation by binding of G-actin to tandem actin-binding WASP-Homology 2 (WH2) domains (Dominguez, 2016). The dynamics of the actin cytoskeleton are also regulated by the actin capping protein (CP), a heterodimer of structurally similar α- and β-subunits. CP binds to the barbed ends of actin filaments and restricts the length of the filaments by preventing filament elongation or dissociation (Edwards et al., 2014). Extensive studies have revealed that CP participates in many cellular processes, including lamellipodia and filopodia formation (Mejillano et al., 2004) and regulation of endosomal trafficking by fine-tuning F-actin density around endosomes (Wang et al., 2021). Importantly, the capping activity of CP is further regulated by multiple proteins that contain a conserved capping protein interaction (CPI) motif. These CPI motif-containing proteins recruit CP to specific cellular membrane locations (Edwards et al., 2015) and/or allosterically inhibit the capping activity of CP (Bruck et al., 2006).

Given the essential role of actin in cell physiology, many bacterial pathogens have evolved distinct strategies to target the host actin cytoskeleton to promote their survival, proliferation, and dissemination (Haglund and Welch, 2011; Stradal and Schelhaas, 2018). Various extracellular pathogens deliver bacterial toxins and effectors to modify Rho family GTPases or actin through ADP-ribosylation, as well as other types of posttranslational modifications to disrupt host actin homeostasis and thus prevent pathogen uptake (Aktories, 2015; Aktories et al., 2011). In contrast, intracellular bacterial pathogens secrete effectors mimicking actin nucleators to promote host actin polymerization and facilitate host cell entry. For example, the virulence factor VopL from *V. parahaemolyticus,* like many other eukaryotic nucleators, dimerizes and promotes actin nucleation via its tandem WH2 domains (Bugalhao et al., 2015; Dominguez, 2016; Namgoong et al., 2011).

Recent studies have revealed that several *L. pneumophila* effectors target the host actin cytoskeleton. VipA is found as an actin nucleator with an unknown mechanism (Bugalhao et al., 2016; Franco et al., 2012). By altering the host actin cytoskeleton, VipA interferes with host membrane trafficking and promotes the invasion of epithelial cells by filamentous *Legionella pneumophila* (Franco et al., 2012; Prashar et al., 2018). The *L. pneumophila* effector, RavK is reported to disrupt actin structures by direct proteolytic cleavage of actin (Liu et al., 2017). Two other *Legionella* effectors, LegK2 and WipA, target the Arp2/3 complex by phosphorylation or dephosphorylation modifications, respectively, to inhibit actin polymerization (He et al., 2019; Michard et al., 2015). Despite accumulating evidence, the exact mechanism and the biological significance of actin hijacking during *Legionella* infection are largely unknown.

In a screen to search for *Legionella* effectors that perturb host actin dynamics, we identified the *L. pneumophila* effector MavH localizes to endosomes and promotes actin polymerization on the surface of the endosome. We found that the intact C-terminal PI(3)P (phosphatidylinositol-3 phosphate) binding domain of MavH is required for its endosomal localization and actin patch formation. We further showed that MavH has a CPI motif that recruits CP to the endosome. Our *in vitro* actin polymerization assays revealed that MavH inhibits actin polymerization in solution, however, it promotes robust actin polymerization in the presence of PI(3)P-containing liposomes. Interestingly, we showed that MavH localizes to the surface of the LCV at the very early stage of infection and correlates with F-actin signals. Together, our results reveal a novel mechanism of actin polymerization, which is catalyzed by a single WH2 domain protein in a PI(3)P-containing membrane-dependent manner.

## RESULTS

### MavH induces F-actin patches around the endosome

Since actin is conserved in all eukaryotes and is an essential target by a variety of pathogens, we performed a screen for *Legionella* effectors that perturb the host actin cytoskeleton. In this screen, 315 *Legionella* effectors were fused with an N-terminal GFP tag and transfected in HeLa cells. The cells were then stained with phalloidin for F-actin. In this screen, we identified MavH as a potential candidate that causes actin rearrangement. GFP-MavH exhibited a punctate localization and colocalized with the early endosomal marker EEA1 (Figure. 1A) and the PI(3)P marker RFP-FYVE (Figure 1—figure supplement 1) when exogenously expressed in HeLa cells. Interestingly, strong F-actin signals were observed on MavH-positive endosomes (Figure. 1B), indicating that MavH may cause actin rearrangement on the surface of endosomes. Moreover, exogenous expression of MavH also causes endosomal trafficking defects as evidenced by the delayed trafficking of EGF in cells transfected with MavH (Figure 1—figure supplement 2).

**Figure 1.**
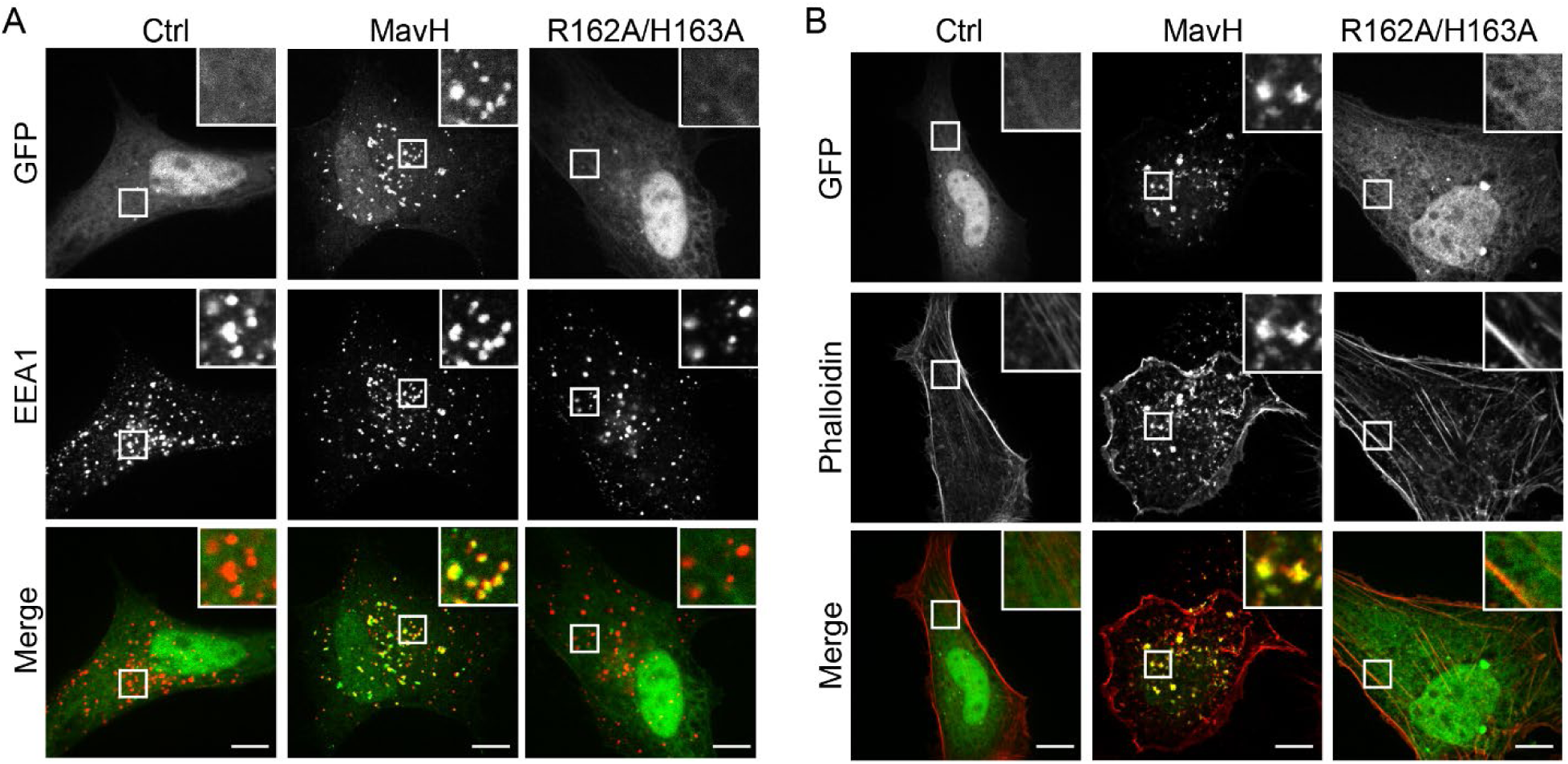
MavH causes actin patch formation around endosomes. (A) Localization of MavH in mammalian cells. HeLa cells were transfected with a plasmid expressing either GFP, GFP-MavH, or GFP-MavH R162A/H163A for 20 hours. Cells were fixed and immuno-stained with EEA1 antibodies and imaged by confocal microscopy. (B) Effect of MavH on the actin cytoskeleton in mammalian cells. HeLa cells were transfected with a plasmid expressing either GFP, GFP-MavH, or GFP-MavH R162A/H163A for 20 hours. Cells were then fixed and stained with Rhodamine conjugated phalloidin and imaged by confocal. Scale bars, 10 *µ*m.

MavH was previously shown to interact with PI(3)P lipids via its C-terminal lipid binding domain (Nachmias et al., 2019). Structure prediction with AlphaFold2 revealed that the C-terminal domain (CTD) of MavH has a compact all-alpha-helical fold (Jumper et al., 2021) (Figure 1—figure supplement 3A and B). We first characterized the lipid-binding specificity by MavH. We performed liposome co-sedimentation assays and revealed that MavH binds preferentially to PI(3) P-containing liposomes (Figure 1—figure supplement 3C and D). We next mapped the key residues involved in PI(3)P binding. According to the structure, a pocket with positive electrostatic surface potentials is evident on the surface of the CTD, which is predicted to mediate PI(3)P binding (Figure 1—figure supplement 3E and F). Indeed, PI(3)P-binding was substantially impaired by the MavH R162A/H163A mutant, which was designed to disrupt the positive charges at the predicted PI(3)P binding pocket (Figure 1—figure supplement 3G and H). In agreement with the *in vitro* assays, MavH R162A/H163A mutant exhibited a cytosolic localization and no endosomal actin patches were observed in cells expressing this mutant (Figures 1A and B). Together, these results suggest that the PI(3)P-binding CTD is required for MavH endosomal localization and actin patch formation around the MavH-positive endosomes.

### MavH interacts with Capping Protein (CP) via a conserved CPI motif

To elucidate the molecular mechanism of MavH in actin polymerization, we performed a sequence analysis of MavH using HHpred (https://toolkit.tuebingen.mpg.de/tools/hhpred). We found that the central region of MavH contains a conserved sequence stretch that resembles the capping protein interaction (CPI) motif, which has a consensus sequence as LxHxTxxRPK(6x)P (Figure 2A). The CPI peptide wraps around the stalk region of the mushroom-shaped CP complex and targets the CP to specific cellular membrane locations to regulate the dynamics of the actin cytoskeleton (Edwards et al., 2015; Hernandez-Valladares et al., 2010). To test whether MavH has a functional CPI motif, we first co-expressed the CP complex (HA-tagged α subunit and mCherry-tagged β subunit) with GFP-MavH in HEK-293T cells to analyze the recruitment of CP by MavH. CP showed a diffused cytosolic localization in cells expressing GFP control whereas it colocalized with MavH to punctate structures in cells expressing wild-type GFP-MavH. Interestingly, the MavH CPI motif mutant, GFP-MavH R73A/K75A failed to recruit CP to punctate structures when overexpressed in cells (Figure 2B). Furthermore, the colocalization between MavH and CP to punctate structures was detected in cells expressing a MavH truncation mutant (MavH Δ53), of which the N-terminal 53 residues were deleted, but not in cells expressing the MavHΔ93 mutant, which lacks the CPI motif (Figure 2B). Next, we assessed the direct interaction between MavH and CP. GFP-tagged MavH and the CP complex were co-transfected in HEK-293T cells and cell lysates were prepared after transfection for 2 days. GFP-MavH was immunoprecipitated by the resin conjugated with anti-GFP nanobodies and the CP α subunit was detected from materials co-immunoprecipitated with wild-type MavH but not its CPI mutant (MavH R73A/K75A) (Figure 2C). The interaction between MavH and CP appears to be direct as evidenced by the pull-down of purified CP complex with the immobilized recombinant MavH protein (Figure 2D). These results suggest that the MavH CPI motif is required to mediate the interaction with CP and intracellular recruitment of CP to punctate structures.

**Figure 2.**
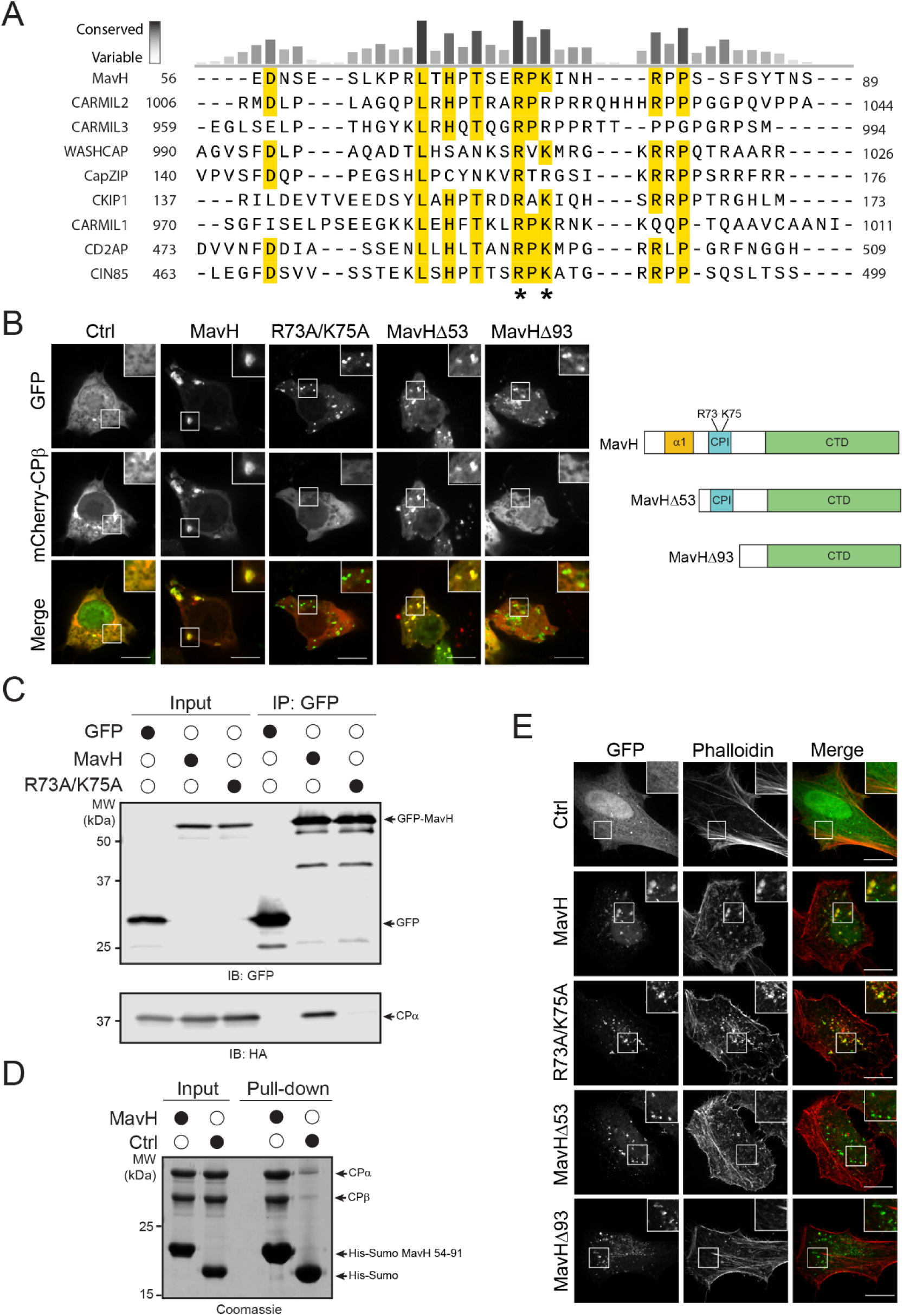
MavH recruits Capping Protein (CP) via a conserved CPI motif. (A) Multiple sequence alignment of MavH with the CPI motif family. The sequences corresponding to the CPI motif were aligned by Clustal Omega. Identical residues and similar residues are highlighted in yellow. Two conserved positive charged residues (R73 and K75) are highlighted with “*”. Uniprot accession numbers for MavH: Q5ZSU1; CARMIL2: Q6F5E8; CARMIL3: Q8ND23; WASHCAP: Q9Y4E1; CapZIP: Q6JBY9; CKIP1: Q53GL0; CARMIL1: Q5VZK9-1; CD2AP: Q9Y5K6; CIN85: Q96B97. (B) Recruitment of CP by MavH is dependent on the CPI motif mutant. GFP-tagged MavH constructs were co-expressed with mCherry-CP*β*-HA-CP*α* in HEK293T cells. Cells were fixed and imaged by confocal. Scale bars, 10 *µ*m. (C) Co-immunoprecipitation of GFP-tagged MavH proteins with CP. HEK293T cells were co-transfected with mCherry-CPβ-HA-CPα with GFP empty vector or GFP-tagged MavH or GFP-tagged MavH R73A/K75A. GFP-tagged proteins were immunoprecipitated from whole-cell lysates with anti-GFP antibodies and then analyzed by SDS-PAGE followed by immunoblot with both anti-HA and anti-GFP antibodies. (D) In vitro pull-down of CP by His-Sumo or His-Sumo-tagged MavH. Purified His-Sumo or His-Sumo-tagged MavH (a.a. 54-91) as bait proteins were loaded onto cobalt beads to pull down recombinant CP. Pull-down materials were resolved by SDS-PAGE and stained by Coomassie-blue dye. (E) The rearrangement of the actin cytoskeleton around endosomes by MavH is independent of the CPI motif. Hela cells were transfected with GFP tagged MavH, CPI motif mutant or truncations, and stained actin with Rhodamine conjugated phalloidin. Scale bars, 10 *µ*m.

We then asked whether the CPI motif of MavH is also responsible for the actin patch formation at endosomes. To test this, we transfected GFP-tagged MavH wild type and mutants into HeLa cells and stained the cells with phalloidin. Interestingly, the CPI motif mutant, MavH R73A/K75A still induced actin patches around the endosome comparable to wild-type MavH (Figure 2E). However, no significant actin patch was detected in cells expressing MavH Δ93 or even MavH Δ53, which has an intact CPI motif and can recruit the CP to endosomes (Figure 2E). These results suggest that the N-terminal region, but not the CPI motif, is responsible for actin recruitment and polymerization at the endosome.

### MavH contains an N-terminal WH2-like domain and interacts with actin

To address how MavH promotes actin assembly, we analyzed the primary sequence of the N-terminal region of MavH. Multiple sequence alignment revealed that several conserved hydrophobic residues form a cluster at the beginning of the predicted N-terminal α helix (Figures 3A and B). This structural feature is reminiscent of the actin-binding WH2 domain, which consists of one α helix with few exposed hydrophobic residues engaging in a hydrophobic cleft formed between the subdomains 1 and 3 on actin (Dominguez, 2004). To test whether the N-terminal α helix mediates the interaction with actin, we expressed GFP-tagged MavH and its mutants in HEK293T cells and assessed the interaction between MavH and actin by co-immunoprecipitation. Indeed, wild-type MavH, as well as its PI(3)P-binding mutant, was able to pull down actin, while the interaction with actin was substantially impaired by the MavH-V24D/L31D mutant, of which the two most conserved N-terminal hydrophobic residues were mutated (Figure 3C). The interaction between actin and MavH was further mapped to the N-terminal region containing the predicted first α helix, as evidenced by the pull-down of actin by MavH 13-65 but not the MavH 13-65 V24D/L31D mutant (Figure 3C). These data suggest that MavH harbors an N-terminal α helix that resembles a WH2 domain and mediates actin binding.

**Figure 3.**
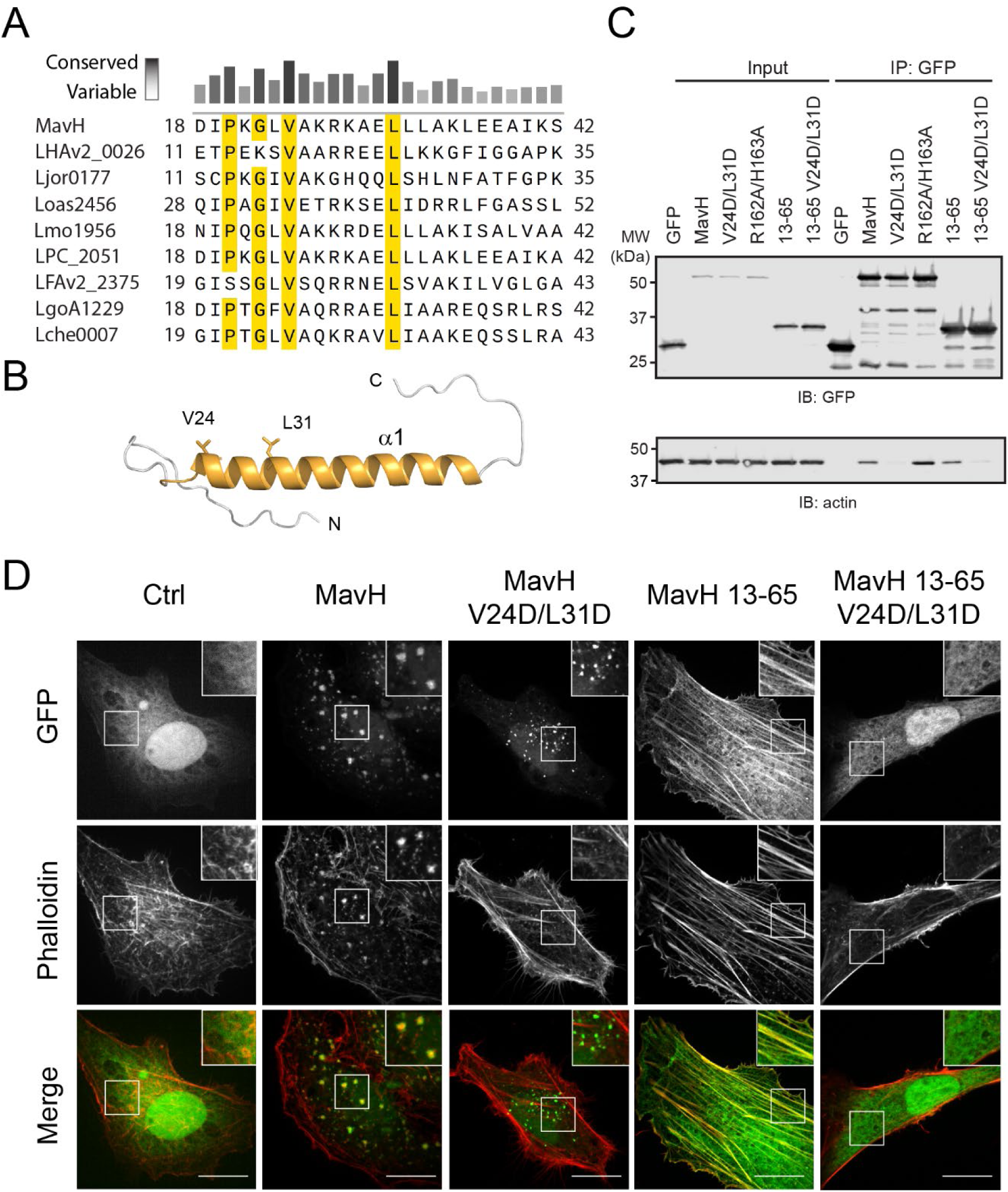
MavH contains an N-terminal WH2-like domain and interacts with actin. (A) Multiple sequence alignment of N-terminus of MavH with other *Legionella* homologs. Identical residues and similar residues are highlighted in yellow. Of note, highly conserved residues are all hydrophobic. (B) Predicted structure of MavH N terminus with AlphaFold2. Two highly conserved hydrophobic residues (V24D and L31D) are shown in sticks. (C) MavH interacts with actin via the N terminus. HEK293T cells were transfected with GFP empty vector or GFP tagged MavH or GFP tagged MavH truncations or mutants. GFP-tagged proteins were immunoprecipitated from whole-cell lysates with anti-GFP antibodies and then analyzed by immunoblot with both anti-actin and anti-GFP antibodies. (D) The rearrangement of the actin cytoskeleton caused by MavH is dependent on interaction with actin via the N terminus. Hela cells were transfected with GFP tagged wild type MavH, mutants, or truncations and stained actin with Rhodamine conjugated phalloidin. Scale bars, 10 *µ*m.

To investigate the effects of the N-terminal WH2-like domain of MavH on actin dynamics, we overexpressed GFP-tagged MavH and its mutants in HeLa cells and examined their localization and actin structures in transfected cells. Strikingly, although MavH-V24D/L31D showed a punctate localization, no actin signals were detected on the MavH positive puncta in contrast to wild-type MavH (Figure 3D). In agreement with the co-IP assay, the N-terminal WH2-like domain alone was sufficient to target GFP-MavH-WH2 to the actin cytoskeleton, whereas the V24D/L31D mutant counterpart was completely cytosolic (Figure 3D).

To further validate the role of the N-terminal WH2-like domain, we fused the N-terminal region of MavH with another PI(3)P binding domain from a *Legionella* effector SetA, which is localized to endosomes when expressed in eukaryotic cells (Beck et al., 2022; Beck et al., 2020). Similar to wild-type MavH, the fusion protein also exhibits an endosomal localization and induces actin patch formation around the endosomes (Figure 3—figure supplement 1). These results support that the N-terminal portion of MavH is responsible for actin polymerization.

MavH intracellular localization and MavH-mediated actin polymerization on intracellular membrane-bound organelles can be further recapitulated in yeast (Figure 3—figure supplement 2). We transformed GFP or mCherry-tagged MavH constructs in yeast cells that were under the control of a galactose inducible promoter. Wild-type MavH, as well as MavH constructs that have an intact CTD, was found to be enriched on the surface of yeast vacuoles. In contrast, the PI(3)P-binding mutant (MavH-R162A/H163A) and the N-terminal region of MavH (MavH-WH2) showed a peripheral punctate localization (Figure 3—figure supplement 2A). These results suggest that the localization of MavH to the vacuole is dependent on its C-terminal PI(3)P-binding domain. Like in mammalian cells, wild-type MavH and its CPI mutant (MavH-R162A/H163A) induced robust actin polymerization on the surface of the vacuole. However, MavH mutants are either incapable of actin-binding (MavH V24D/L31D and MavH CTD) or defective in membrane binding (MavH-R162A/H163A and MavH-WH2) failed to polymerize actin on the vacuole (Figure 3—figure supplement 2B). Furthermore, MavH-R162A/H163A and MavH-WH2 displayed colocalization with peripheral actin patches, consistent with the binding of MavH WH2-like domain with actin. Interestingly, over-producing MavH constructs that contain the intact WH2-like domain are toxic in yeast (Figure 3—figure supplement 2C), indicating that the WH2-like domain may interfere with endogenous actin dynamics and causes yeast growth defects.

In summary, our results revealed that the N-terminal region of MavH harbors an actin-interacting motif. This WH2-like domain, together with its C-terminal PI(3)P-binding domain, is responsible for actin assembly on the surface of endosomes.

### Membrane-dependent actin polymerization mediated by MavH

To elucidate the molecular mechanism of actin assembly catalyzed by MavH, we performed in vitro pyrene-actin polymerization assays (Harris and Higgs, 2006). To our surprise, wild-type MavH did not promote actin assembly in F-actin buffer, instead, it inhibited actin polymerization compared to the actin alone control (Figure 4A). A similar inhibitory effect was also observed for the lipid-binding motif mutant (R162A/H163A). However, the MavH WH2 mutant (V24D/L31D) showed no effect on actin polymerization (Figure 4A). These results suggest that MavH binds to actin via its single WH2-like domain and this binding sequesters actin from polymerization in solution. Since MavH promoted actin assembly on endosomes in the cell, we reasoned that MavH-triggered actin assembly may require the membrane. To test this idea, we performed the pyrene-actin assay in the presence of PI(3)P-containing liposomes (PC : PS : PI(3)P = 8 : 1 : 1). Liposomes alone did not affect actin polymerization, however, in the presence of both liposomes and wild type MavH, actin polymerization was enhanced, particularly at the initial stage (Figure 4B). Surprisingly, the fluorescence signals exhibited an abnormal fluctuating reading. To explain the unexpected reading, we performed similar actin polymerization assays and visualized the final products by confocal microscopy following the staining with 488-phalloidin. Strikingly, massive F-actin was observed congregating around the liposomes, concomitant with liposome deformation and clustering (Figure 4C). TEM analysis further revealed that membrane tubules were induced from deformed liposomes by MavH-mediated actin polymerization and membrane tubules were decorated with longitudinal F-actin fibers (Figure 4D). As a control, the MavH-V24D/L31D showed no effect on actin polymerization (Figure 4B), and no significant F-actin signals were observed around the liposomes (Figure 4C).

**Figure 4.**
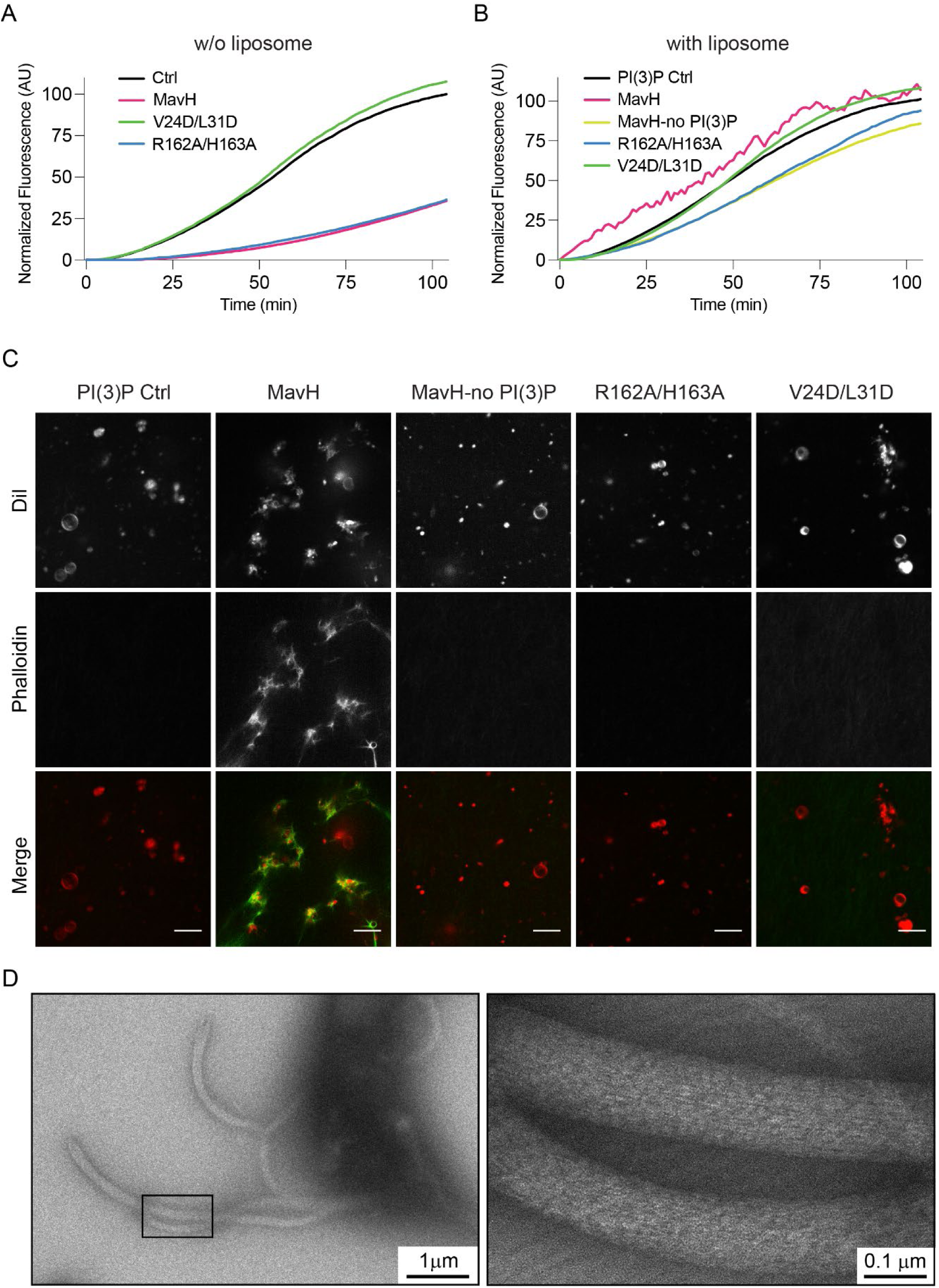
MavH promotes actin polymerization on PI(3)P-positive liposomes. Pyrene-actin polymerization assays of MavH without (A) and with (B) PI(3)P-containing liposomes. All reactions contain 3 *µ*M actin (10% Pyrene-actin) and 250 nM wild-type or mutant MavH proteins. Actin polymerization was initiated by adding 10X actin polymerization buffer and fluorescence (AU) signals were recorded over time. (C) In vitro liposome imaging assay. Reactions were performed using 3 µM actin, 250 nM MavH. Liposomes were used at 500 µM. After induction of actin polymerization, actin was stained with 488-Phalloidin. Images were taken by confocal microscopy. (D) EM images of negatively stained PI(3)P-containing liposomes incubated with actin and MavH after induction of actin polymerization. Images were taken at a magnification of 5,300X (left) and 92,000X (right). Reactions were performed using 6 µM actin, 500 µM PI(3)P-containing liposomes, and 1 µM MavH.

The importance of membrane binding in MavH-mediated actin polymerization was further validated by in vitro pyrene-actin assays when the membrane association of MavH was perturbed. The PI(3)P-binding defective mutant, MavH-R162A/H163A showed no stimulation of actin polymerization (Figure 4B), and no F-actin was detected around the liposomes (Figure 4C). Along this line, liposomes lacking PI(3)P also failed to promote actin polymerization triggered by wild-type MavH (Figures 4B and C). Together, these data suggest that although MavH contains a single actin-binding WH2-like domain and inhibits actin polymerization in solution, however, it promotes F-actin assembly on PI(3)P-containing membranes upon its association with the membrane via its C-terminal PI(3)P-binding domain.

### MavH recruits CP to negatively regulate actin polymerization at the membrane

We next asked about the role of the CPI motif in MavH-mediated actin assembly. It has been reported that capping protein regulates F-actin density around endocytic vesicles (Durre et al., 2018; Wang et al., 2021). We speculate that MavH might recruit CP to regulate F-actin density at PI(3)P-containing membranes. To test this hypothesis, we first generated a CP mutant (CPβ-R15A), which carries an R15A substitution at its β subunit. This mutant is defective in binding with the CPI motif (Edwards et al., 2015) and hence the recruitment to the liposome by MavH (Figures 5A and B). However, it maintains a comparable capping activity as wild-type CP (Figure 5C). We then used pyrene-actin assay to analyze the effect of the wild type and the mutant CP on MavH-catalyzed actin polymerization. We observed that wild-type CP substantially inhibited MavH-mediated actin polymerization while the CPβ-R15A mutant displayed a milder inhibition on actin polymerization (Figure 5D). Correspondingly, the F-actin signal around the liposome was substantially weaker and the liposomes were less aggregated in the presence of wild-type CP compared to that of CPβ-R15A mutant (Figure 5E). Together, these data suggest that the recruitment of CP via the CPI motif negatively regulates actin polymerization.

**Figure 5.**
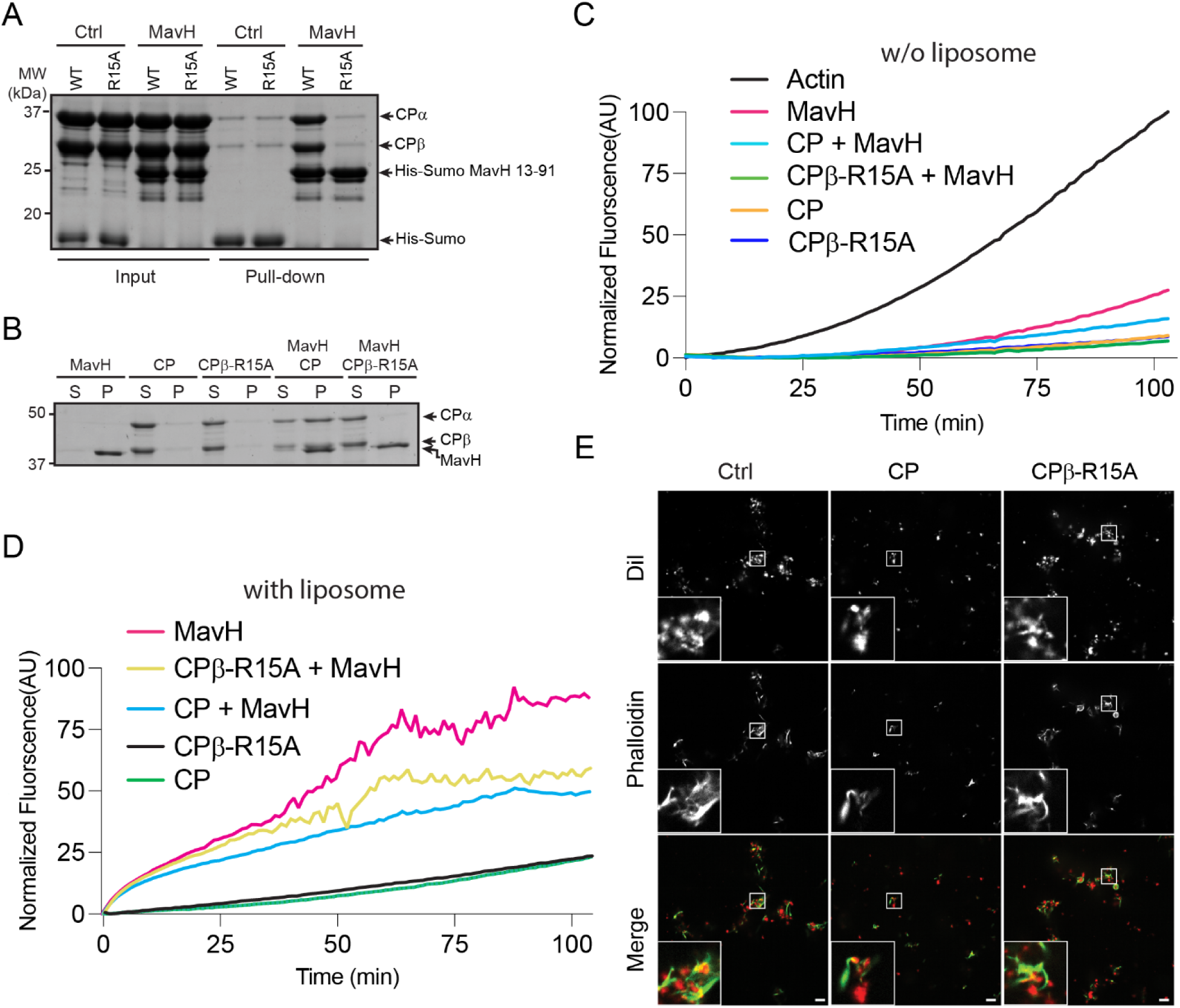
The CPI motif of MavH recruits CP and negatively regulates actin polymerization. (A) In vitro pull-down assay of CP by MavH. Cobalt beads preloaded with His-Sumo or His-Sumo-tagged MavH (a.a. 13-91) were used as the bait to pull down purified wild-type CP or CP*β*-R15A. Pull-down materials were resolved by SDS-PAGE and stained by Coomassie-blue dye. (B) Liposome co-sedimentation assay of CP or CPβ-R15A with MavH. Purified proteins were incubated with PI(3)P-containing liposomes and then spun down by ultracentrifugation. The pellets were analyzed by SDS-PAGE followed by Coomassie-blue staining. (C) Pyrene actin polymerization assay with actin alone or in the presence of CP, CP*β*-R15A, MavH. (D) Pyrene actin polymerization assay of CP, CP*β*-R15A and MavH in the presence of PI(3)P containing liposomes. (E) In vitro liposome imaging assay. Reactions were performed using 3 µM actin, 250 nM MavH, 25 nM CP, or CPβ-R15A mutant. PI(3)P-containing liposomes were used at 250 µM. After 30 min incubation, actin was stained with 488-Phalloidin, and the reaction products were imaged with fluorescence confocal microscopy. Scale bars, 10 *µ*m.

### MavH localizes to the LCV membrane at the early stage of *Legionella* infection

We next examined the intracellular localization of MavH during intracellular infection by *L. pneumophila*. We first created a MavH deletion strain and strains supplemented with a plasmid expressing wild-type or mutant MavH fused with an N-terminal 4xHA tag. These strains were then used to infect HEK293T cells expressing FcγRII receptor. After infection for 10 min, cells were fixed with ice-cold methanol and immunostained with an anti-HA antibody. HA signals were detected around the LCV in cells infected with Lp02Δ*mavH* supplemented with 4xHA-MavH but not with Lp03 overexpressing 4xHA-MavH (Figure 6A). We then inspected the time course of the retention of MavH at the LCV. MavH was detected at the LCV as early as 2 min p.i. and peaked at around 5 min p.i. (~ 20% LCVs are positive for MavH). MavH signals were gradually reduced as the infection progressed (Figure 6B). We further investigated the functional determinants for the anchoring of MavH to the LCV. We observed that the actin-binding mutant, MavH-V24D/L31D showed a slight reduction of LCV localization while the CPI mutant, MavH-R73A/K75A exhibited no discernable difference compared to the wild type. However, nearly no HA signals could be detected at the LCV for the PI(3)P-binding mutant, MavH-R162A/H163A (Figures 6C and D). We next investigated the role of MavH in *L. pneumophila* intracellular proliferation. The intracellular growth of the MavH deletion strain, as well as strains that were supplemented with a plasmid expressing either wild-type 4xHA-MavH or MavH mutants, showed no obvious defects in *Acanthamoebae castellanii* compared to that of the wild-type strain (Figure 6E). Together, these results demonstrated that MavH localizes to the LCV at the early stage of *Legionella* infection and the anchoring of MavH to the LCV requires its C-terminal PI(3)P-binding domain. Although MavH is dispensable for intracellular growth in *A. castellanii,* it may play a role in the early stage of bacterial infection.

**Figure 6.**
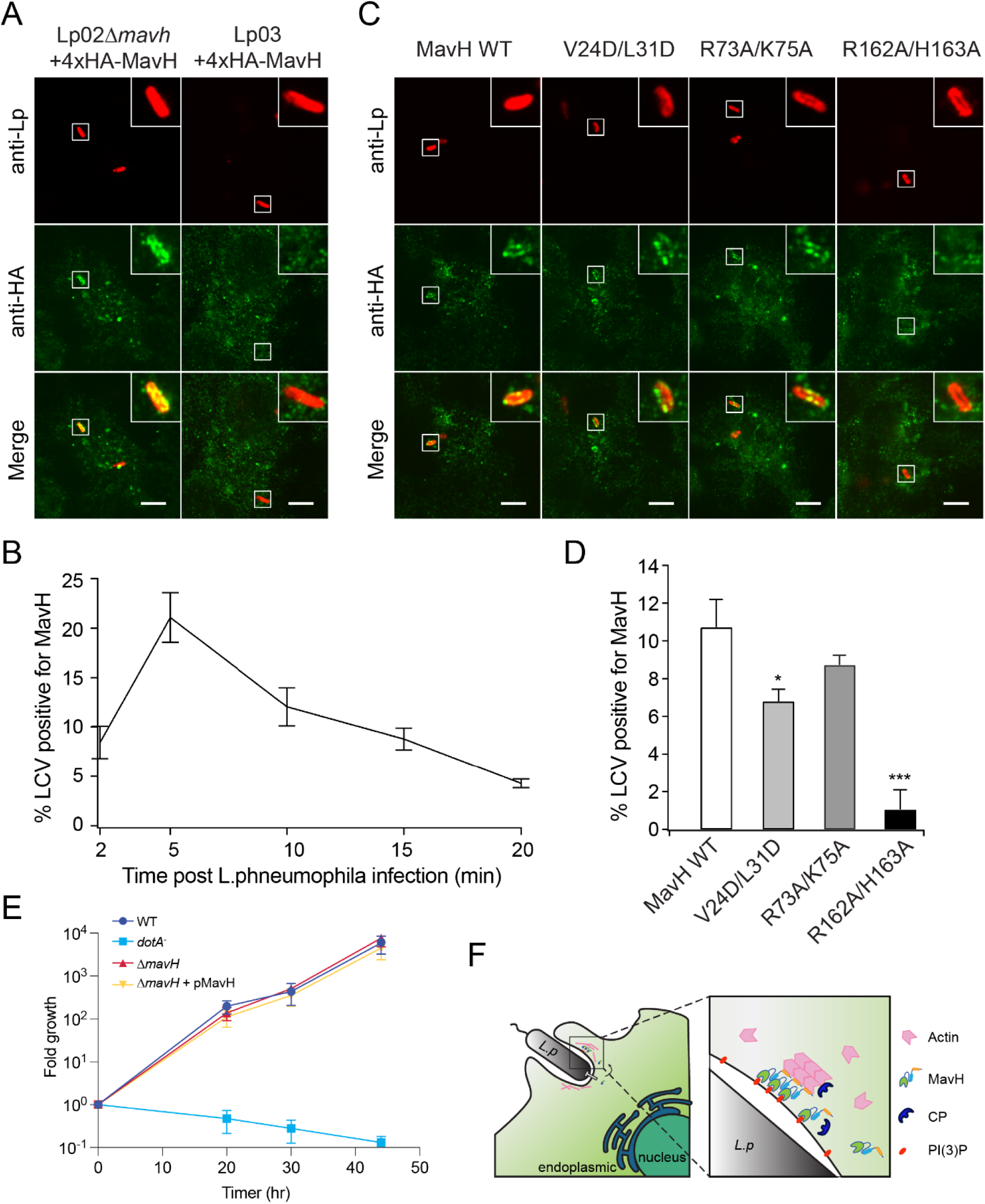
MavH localizes to the surface of LCV at the early stage of infection. (A) FcγRII-expressing HEK293T were challenged by *mavH* deletion Lp02 or Lp03 strains supplemented with a plasmid expressing 4xHA-MavH for the indicated time. Cells were fixed using 4% PFA, for 15 min and then permeablized using ice-cold methanol for 10 min. 4xHA-MavH was immunostained using mouse-anti-HA primary antibodies and Alexa-488 anti-mouse secondary antibodies. Representative images showed localization of 4xHA-MavH at 5 min post-infection. Scale bars = 10 *µ*m. (B) Quantifications of LCVs positive for 4xHA-MavH in HEK293T cells infected by Lp02*ΔmavH* overexpressing 4xHA-MavH for the indicated time, shown as Mean ± SEM from three independent experiments. At least 50 cells were analyzed for each time point. (C) FcγRII-expressing HEK293T cells were infected by *mavH* deletion strain overexpressing 4xHA-MavH wild type or indicated mutant for 10 min. cells were fixed and immunostained with anti-HA antibodies as in (A). Scale bars = 10 *µ*m. (D) Quantifications of MavH-positive LCVs. The percentage of MavH-positive LCVs was shown as Mean ± SEM from three independent experiments. At least 40 cells were analyzed for condition. *P<0.05 and ***P<0.001. (E) Intracellular growth assay of *Legionella* in *A. castellanii* host. A wild-type *Legionella* strain, the Dot/Icm deficient Δ*dotA^-^*, the *mavH* deletion strain, and *mavH* deletion strain overexpressing 4xHA-MavH were used to infect *A. castellanii* cells. Growth was assayed by plating colony-forming units (CFUs) at the indicated time after infection. The growth assays were performed in triplicate. (F) A hypothetic model of MavH at the early stage of *Legionella* infection. MavH is secreted at the early stage of infection to promote actin assembly facilitating the bacterial entry of host cells.

## DISCUSSION

Actin polymerization requires actin nucleation factors to overcome the kinetic barrier and assemble an initial nucleus for elongation by the addition of actin monomers. To date, three major classes of eukaryotic nucleators have been identified: the Arp2/3 complex, the formins, and the tandem actin-binding domain proteins. These actin nucleators apply distinct mechanisms for actin nucleation. The Arp2/3 complex is a seven-subunit complex, of which, the Arp2 and Arp3 subunits form a structural mimic of an actin dimer and serve as the nucleator for F-actin assembly. Upon activation by nucleation-promoting factors, the Arp2/3 complex facilitates actin assembly to form a branched actin filament from an existing actin filament (Goley and Welch, 2006) or linear actin filaments in the absence of a preformed actin filament (Shaaban et al., 2020; Wagner et al., 2013). The formins possess characteristic formin-homology 1 domain, which recruits profilin-actin, and formin-homology 2 domain, which mediates the dimerization of formins and facilitates the addition of actin monomers from profilin-actin to the barbed end of the actin filament (Breitsprecher and Goode, 2013). The third family of actin nucleators, including Spire (Kerkhoff, 2006) and Cobl (Ahuja et al., 2007), contain tandem repeats of actin-binding motifs, such as the WH2 domain. These tandem actin-binding domains serve as a scaffold to recruit actin monomers and synergize with other functional domains for F-actin assembly (Dominguez, 2016). Interestingly, many bacterial actin nucleators are found to fall into one of the three categories. For example, the Vibrio Cholerae virulent effectors, VopL and VopF, mimic the tandem WH2 domain-containing nucleators (Burke et al., 2017; Zahm et al., 2013), while the Rickettsia effector, Sca2 promotes actin assembly like the formins (Madasu et al., 2013). Here we report a novel mechanism of actin polymerization catalyzed by the *Legionella* effector, MavH. Unlike other actin assembly factors, MavH harbors a single actin-binding WH2-like domain and inhibits actin polymerization in solution. However, it promotes robust actin polymerization on the membrane surface upon its binding to PI(3)P-containing liposomes or membrane-bound organelles. Moreover, MavH-mediated actin polymerization also triggers membrane tubulation and the membrane tubules are likely stabilized by longitudinally bound F-actin fibers. These observations raise intriguing questions, for example, how does MavH facilitate actin polymerization on membrane surface; and how does MavH-mediated actin polymerization induce membrane deformation and tubulation? Future experiments, such as high-resolution Cryo-EM studies, are needed to address the molecular mechanism of MavH-mediated actin polymerization. Nevertheless, our results uncover a novel factor that promotes actin assembly in a membrane-dependent manner. Our results may also inspire the discovery of new MavH-like actin polymerization factors in other pathogens or eukaryotes.

MavH is a unique actin assembly promotor in that it contains a CPI motif following the actin-binding WH2-like domain. In this study, we showed that MavH recruits CP to endosomes when ectopically expressed in mammalian cells. Moreover, we found that MavH can modulate actin dynamics and actin density around the liposomes through its recruitment of CP via its CPI motif. Nevertheless, the physiological consequences of CP recruitment by MavH during infection are not known. CPI motif-containing proteins have been shown to recruit CP to specific cellular locations (Edwards et al., 2015) and/or regulate actin-capping activity by allosteric effects (Bruck et al., 2006; Hernandez-Valladares et al., 2010). Aside from terminating filament growth, a recent study showed that capping the barbed ends of actin filaments facilitates branched actin network assembly (Funk et al., 2021). Thus, we speculate that MavH may fine-tune actin dynamics and possibly promote branched actin network formation, which is important for cellular membrane movement in a number of cellular processes, including phagocytosis. However, further experiments are needed to elucidate the biological significance of the recruitment of CP in MavH-mediated actin assembly.

Dynamic remodeling of the actin cytoskeleton is essential for cell physiology. Many intracellular bacterial pathogens have involved distinct strategies to alter host actin cytoskeleton dynamics at different infection stages, including entry into host cells (Dramsi and Cossart, 1998; Rottner et al., 2005), actin-based intracellular movement (Dramsi and Cossart, 1998; Lamason and Welch, 2017), evasion of endocytic degradation by the formation of the “actin cocoon structure” around the bacterial containing vacuole (Kuhn et al., 2020). It is interesting to note that several other *Legionella* effectors were found to perturb the dynamics of the host actin cytoskeleton. The *Legionella* effector VipA was shown to promote F-actin assembly and alters host cell membrane trafficking, however, the mechanism for promoting actin polymerization by VipA was not understood (Bugalhao et al., 2016; Franco et al., 2012). Another *L. pneumophila* effector, RavK is reported to disrupt actin structures by direct proteolytic cleavage of actin (Liu et al., 2017). Two other *Legionella* effectors, LegK2 and WipA, alter the phosphorylation state of the host Arp2/3 complex and inhibit action polymerization (He et al., 2019; Michard et al., 2015). These studies underscore the importance of actin in *Legionella* infection and shed light on the intricate control of the host actin cytoskeleton during the infection process. In this study, we identified a novel *L. pneumophila* actin polymerization promotor that triggers actin polymerization in a membrane-dependent manner. We also found that MavH was delivered at the very early stage of infection and localized to the LCV via its binding to PI(3)P. These observations led us to hypothesize that MavH may drive actin polymerization and membrane deformation at the phagocytic site to facilitate the uptake of the bacterium (Figure 6F). It is interesting for future studies to elucidate how these effectors, which have synergic or antagonistic activities on actin dynamics, orchestrate to exploit the host actin cytoskeleton for successful infection.

## MATERIALS AND METHODS

### Cloning and Site-Directed Mutagenesis

Full-length MavH (a.a. 1-266) was amplified from *L. pneumophila* genomic DNA and digested with BamHI/SalI and inserted into a pET28a-based vector in-frame with an N-terminal His-SUMO tag. PCR products for MavH truncations were amplified from the constructed pET28a His-Sumo MavH. Mutations of MavH were introduced by in vitro site-directed mutagenesis using specific primers containing the defined base changes and PrimeSTAR® Max DNA Polymerase (Takara Bio, Inc.) premix. For mammalian expression, corresponding fragments of MavH were subcloned into pEGFP-C1 vector. For the MavH-SetA chimeric fusion, the N-terminus of MavH (a.a. 13-65) was amplified and digested with BamHI and SalI and then ligated into the pEGFP-C1 vector digested with BglII and SalI. The C-teriminal PI(3)P-binding domain of SetA (a.a. 507-629) was amplified and digested with SalI and BamHI and then ligated into pEGFP-MavH (a.a. 13-65) digested with SalI and BamHI to generate pEGFP-MavH-SetA for the expression of the MavH-SetA fusion protein.

Bacterial expression of CP plasmid was purchased from Addgene (Plasmid #89950). The α subunit of CP was subcloned into a pRSFDuet-based vector in frame with an N terminal His-Sumo tag and the β subunit of CP was subcloned into a pCDFuet-based vector ORF2. A single residue mutation of CPβ-R15A was introduced by site-directed mutagenesis. For mammalian expression of CP, we constructed pCW57-mCherry-CPβ-P2A-HA-CPα. First, HA-tagged CP alpha subunit was amplified and digested with AvrII and BamHI and cloned into pCW57-P2A vector to generate pCW57-P2A-HA-CPα. Then mCherry-CPβ was first cloned into the mCherry-C1 vector (restriction sites BglII and SalI) and mCherry CPβ was then amplified and digested with NheI and SalI and inserted into pCW57-P2A-HA-CPα to finally generate pCW57-mCherry-CPβ-P2A-HA-CPα.

For *Legionella* expression, MavH was subcloned into a pZL507-based vector (gift from Dr. Zhao-Qing Luo, Purdue University) with 4xHA tag. For yeast expression, corresponding fragments of MavH were subcloned into p415gal-yemCherry and p415gal-yeGFP vectors (gift from Dr. Anthony Bretscher, Cornell Univerisity). All constructs were verified by DNA sequencing.

### Protein Expression and Purification

All MavH constructs in pET28a-His-Sumo were transformed into the Rosetta (DE3) strain of *E.coli* cells using the antibiotic selection markers kanamycin and chloramphenicol. For CP proteins expression, pRSFDuet HisSumo-CPα subunit and pCDFDuet CPβ subunit were co-transformed into the Rosetta (DE3) strain of *E.coli* cells using the antibiotic selection markers kanamycin and spectinomycin. Transformed bacterial cells were grown in 1L expression cultures at 37 °C at 220 rpm and induced with 0.2 mM IPTG during log-phase growth (O.D._600_ = 0.6-0.8). Cells were incubated at 18 °C and 180 rpm for 18 hours post-induction. Cells were collected by pelleting expression cultures at 4000 rpm for 30 minutes at 4 °C. Cells were resuspended in 35 mL of 20 mM Tris (pH 7.5) and 150 mM NaCl containing 1 mM PMSF. Cells were lysed by two rounds of sonication at 50% amplitude, 2-minute duration, and 2 sec on/off pulse on ice. Sonicated samples were spun at 16,000 rpm for 30 minutes at 4 °C to remove the insoluble fraction. The supernatant was collected and mixed with 2 mL of cobalt resin and incubated while rotating for 2 hours at 4 °C to bind proteins. The protein-bound resin was washed with several column volumes of buffer containing 20 mM Tris (pH 7.5) and 150 mM NaCl to remove unbound and nonspecifically bound proteins. The resin was resuspended in 4 mL of wash buffer and cut overnight with His-tagged Ulp1 at 4 °C. Cut proteins were eluted the next day and concentrated to a final volume of 3 mL using a 30 kDa cut-off centrifugal concentrator. Proteins were run on a Superdex200 16/200 column using an AKTA GE Healthcare FPLC system. Peak fractions were collected and analyzed by SDS-PAGE. Purified proteins were further concentrated and stored at −80 °C.

### Cell culture, transfection, and fluorescent microscopy

Hela, Cos7, RAW 264.7, and HEK293T cells were cultured in Dulbecco’s modified minimum Eagle’s medium (DMEM) supplemented with 10% FBS fetal bovine/calf serum (FBS). For co-localization analysis, EGFP-MavH constructs were co-expressed with RFP-2xFYVE domain constructs in Hela cells. For intracellular localization, EGFP-MavH constructs were expressed in Hela cells, and endosomes were marked by staining of EEA1 via EEA1 rabbit monoclonal primary antibodies. For CP localization, GFP-tagged MavH constructs were co-expressed with mCherry-CPβ-HA-CPα in HEK293T cells, and 1ug/ml doxycycline was used to induce the expression of CP during transfection.

For imaging, Hela or HEK293T cells were passaged at 25-30% initial density in a 24-well plate in D10 media. Cells were subsequently transfected 24 hours later with 0.15 µg of each plasmid and a 1:5 (m/v) ratio of polyethyleneimine (PEI) in DMEM for a total volume of 50 µL. At 14-16 hours post-transfection, cells were fixed in 4% paraformaldehyde in PBS solution for 20 minutes on ice and then washed three times with PBS. Fixed coverslips were mounted onto glass slides using Fluoromount-G mounting solution. Fixed cells were imaged using a spinning disk confocal microscope (Intelligent Imaging 108 Innovations, Denver, CO) equipped with a spinning disk confocal unit (Yokogawa CSU-X1), an inverted 109 microscope (Leica DMI6000B), a fiber-optic laser light source, a 100× 1.47NA objective lens, 110 and a Hamamatsu ORCA Flash 4.0 v2+ sCMOS camera. Images were acquired and processed using the Slidebook (version 6) software.

### Immunoprecipitation

HEK293T cells were passaged at 25-30% initial density in a 6-well plate in D10 media. Cells were subsequently transfected with 1.8 µg of each plasmid and a 1:5 (m/v) ratio of polyethyleneimine (PEI) in DMEM for a total volume of 200 µL. At 24 hours post-transfection, cells were washed two times with cold PBS and resuspended in 300 µL of IP lysis buffer (1% Triton-X, 0.1% deoxycholate in 50 mM Tris, pH 8.0, 150 mM NaCl, and protease inhibitor cocktail (Roche)). Cells were briefly sonicated at 10% amplitude for 5 seconds (pulse) and centrifuged at 15000 rpm for 15 minutes at 4°C to remove the insoluble fraction. GFP-nanobody conjugated resin was added to the collected supernatant and incubated for 3 hours on a nutating mixer at 4°C to bind GFP-tagged proteins. Resins with bound proteins were washed with 1 mL of cold PBS for a total of 4 washes. Proteins were eluted from the resin by boiling at 95°C for 4 minutes in 25 µL of SDS sample loading buffer containing 2% BME. Immunoblotting of GFP-MavH was performed using a homemade rabbit anti-GFP antibody at a dilution of 1:1000. Actin was probed using a mouse anti-Actin antibody (Proteintech) at a dilution of 1:1000. HA tagged CPα was probed using a mouse anti-HA antibody (Sigma). Probed proteins were detected using donkey anti-rabbit IgG antibody, DyLight 800 (Invitrogen), and donkey anti-mouse IgG antibody, Alexa Fluor 680 (Invitrogen; cat. no. A10038) secondary antibodies. Membranes were scanned using a LI-COR Odyssey CLx Imager. Western Blot images were processed and analyzed using ImageStudio Lite software (version 5.2). The samples were subsequently probed with mouse anti-HA (Sigma), rabbit anti-GFP, or mouse anti-actin antibody (Proteintech).

### Liposome preparation

1-palmitoyl-2-oleoyl-sn-glycero-3-phosphocholine (POPC) and 1-palmitoyl-2-oleoyl-sn-glycero-3-phospho-L-serine (POPS) and di-C16-phosphatidylinositol polyphosphates were purchased from Avanti. Liposomes were prepared with POPC and POPS (8:2 molar ratio) or POPC, POPS, and Phosphatidylinositols (8:1:1 molar ratio). Lipids mixtures dissolved in 90% chloroform and 10% methanol were dried in glass tubes by nitrogen gas in the fume hood and rehydrated into G-actin buffer followed by 1 hr incubation at 37°C for the spontaneous formation of liposomes. The liposomes for pyrene-actin polymerization were filtered 10 times through 0.1 µm diameter polycarbonate membranes (Nucleopore).

### Liposome co-sedimentation assay

Purified proteins (1 uM) were incubated with 0.25 mM of liposomes for 20 min at room temperature and then spun down in a benchtop ultracentrifuge (Thermo Fisher AccuSpin Micro 17R centrifuge) for 15 min at 17,000 g. Resuspended pellets and supernatants were analyzed by SDS-PAGE, and quantified by Image J. The ratio of protein in pellet was calculated by the protein in pellet/(pellet + supernatant).

### Pyrene actin polymerization assay

Actin was purified from muscle acetone powder as described previously (Pardee and Spudich, 1982) followed by gel filtration at low ionic strength to isolate monomeric ATP-G-actin (MacLean-Fletcher and Pollard, 1980). Purified actin was stored in G-actin buffer (2 mM Tris-HCl (pH 8.0) 0.2 mM CaCl_2_, 0.2 mM ATP, and 0.1 mM DTT). Actin polymerization biochem kit (BK003) was purchased from Cytoskeleton Inc. Actin polymerization assays were performed in 200 µl reactions using 96 well Black polystyrene assay plates. Reactions were started by adding actin to a mix of all other components and 10X actin polymerization buffer (500 mM KCl, 20 mM MgCl_2_, 0.05 M guanidine carbonate, and 10 mM ATP). Fluorescence was measured in a Tecan Safire2 fluorescence plate reader using excitation/emission wavelength 350 nm (± 20 nm) / 410 nm (± 20 nm). All actin polymerization reactions were performed using 3 uM actin (10 % Pyrene labeled actin). Liposomes were used at 50 µM.

### Liposome imaging

Near-infrared Dil dye (Invitrogen) was added when making liposomes to aid the visualization of liposomes. 250 nM wild-type or mutant MavH proteins were incubated with 3uM actin and corresponding liposomes 250 µM at room temperature for 30min. Then polymerization was induced by adding 10X polymerization buffer for 30 min. F-actin was stained by 488-phalloidin 20 min before imaging. For imaging, 5 μL of the reaction mixture was added to the chamber created between a cover slip and a glass slide. Fluorescence microscopy images were acquired using a spinning disk confocal microscope (Intelligent Imaging 108 Innovations, Denver, CO) equipped with a spinning disk confocal unit (Yokogawa CSU-X1), an inverted 109 microscope (Leica DMI6000B), a fiber-optic laser light source, a 63X and a 40X objective lenses. Images were acquired and processed using the Slidebook (version 6) software.

### Sample Preparation and Image Acquisition for Negative Stain

PI(3)P-containing liposomes underwent 5 rounds of freeze-thawing process in liquid nitrogen to make small liposomes. Sample was prepared by incubating 1 µM MavH, 6 µM actin and 500 µM liposomes for 30 min, and actin polymerization was induced by adding 10X polymerization buffer for 30min. Carbon coated EM grids (200 mesh, from Electron Microscopy sciences) were glow-discharged using PELCO easiGlow^TM^ Glow Discharge Cleaning system. A sample of 5 µl was applied to the grids, followed by incubation for 1 min and excess sample was absorbed by paper. Then use 2% uranyl acetate to stain for 1min and absorb the excess stain with paper. Negative stain images were obtained using an F200C microscope.

### EGF trafficking assay

COS-7 cells were split and cultured on poly-lysine-coated cover glass in a 24-well plate. Cells were transfected with the plasmid of EGFP or EGFP-tagged MavH for 24 hours. Cells were incubated with 20 ng/mL Alexa 555-EGF (in DMEM) on ice for 20 min, washed using ice-cold DMEM three times, and then incubated at 37 °C in a CO_2_ incubator for the indicated time. After EGF uptake, cells were fixed using 4% PFA, permeabilized with 0.1% saponin. To label early endosomes, cells were immunostained with rabbit-anti-EEA1 primary antibodies and then with Alexa 647 anti-rabbit secondary antibodies.

### Yeast Strains and growth assays

For MavH localization in yeast cells, the SEY6210.1 yeast strain expressing genomically tagged VPH1-mCherry is obtained from Dr. Scott Emr (Cornell Univ.). This strain was transformed with the plasmid pRS416 Gal3, together with the pRS415 plasmid expressing either GFP-MavH wild type and mutants under a galactose inducible promoter and was grown overnight in a complete supplement mixture −Ura-Leu media (Sunrise Science Products) containing 2% glucose at 30°C. Cultures were then centrifuged at 1000 xg and pellets were resuspended in selective media containing 2% galactose and incubated for 4 hours at 30 °C with agitation to induce protein expression. For the effects of MavH on the actin cytoskeleton in yeast cells, the BY4741 strain expressing mCherry-MavH, truncations, mutants, or with vector controls under galactose inducible promoters was grown overnight in a complete supplement mixture −Leu media containing 2% glucose at 30°C. Cultures were then centrifuged at 500 g and pellets were resuspended in selective media containing 2% galactose and incubated for 4 hours at 30 °C with agitation to induce protein expression. In 3 ml cell cultures, 750 µl 20% PFA was added and incubated for 30min to fix cells and then cells were washed three times with PBS. PBS containing 0.2% Triton-X is used to permeabilize cells and 488-phalloidin (Thermofisher, A12379) is used for actin staining. Cells were immobilized on coverslips using concanavalin A and imaged using a spinning disk confocal microscope. Yeast growth assays were performed as described (Xu et al., 2014). Briefly, yeast cultures for each strain were grown to log phase and diluted to OD_600_ = 0.3 and then serially diluted by a factor of 10. 5 µL of each serially diluted sample was either spotted on −Leu plates containing 2% glucose for repression or 2% galactose for expression of proteins, and then incubated for 2-3 days at 30 °C before imaging.

### *Legionella* strains and infection

Strains of *L. pneumophila* used were the wild type Lp02 and the Dot/Icm deficient Lp03 (Berger and Isberg, 1993). MavH deletion strain was created by a two-step allelic exchange strategy as described (Dumenil and Isberg, 2001). Briefly, a 1.2kb DNA fragment from upstream and downstream of MavH was amplified by PCR including the first 15 amino acids and the last 14 amino acids of the gene. The SalI and BamHI restriction sites were introduced into the 5’ and 3’ end of the upstream fragment while the BamHI and SacI sites were introduced into the 5’ and 3’ end of the downstream fragment, respectively. These fragments digested with the proper enzymes are ligated into plasmid pSR47S (59) and digested with SalI and SacI by three-way ligation. Plasmids were introduced into *L. pneumophila* by conjugation with *E.coli* donors as described previously (Roy and Isberg, 1997). Conjugants containing plasmids integrated into bacterial genome were selected on CYET plates containing kanamycin (20 ug/ml) and streptomycin (50 ug/ml). Kanamycin-resistant *L. pneumophila* colonies were plated on CYET plates containing 5% sucrose to allow selection against the plasmid that contains the sacB gene. MavH deletion strains were verified by PCR. MavH complementary strains were generated by transformation using 4x-HA-tagged mavH or its mutants in the pZL507 vector. 4xHA-MavH expression was induced with 0.1 mM IPTG for 30 min before infection.

To detect MavH localization or F-actin after *Legionella* infection, HEK293T cells were transfected with FcγRII for 24 hrs. Bacteria of indicated *Legionella* strains were opsonized with rabbit anti-*Legionella* antibodies (1:500) at 37 °C for 20 min before infection. The HEK293T cells were infected with post-exponential *L. pneumophila* strains at an MOI of 5 for the indicated amount of time. Cells were then fixed using 4% PFA for 15 min. To detect MavH localization in HEK293T cells, fixed cells were permeablized using ice-cold methanol for 10 min. 4xHA-MavH was immunostained using mouse anti-HA primary antibodies (1:1000) and Alexa 488 anti-mouse secondary antibodies.

### Bacterial growth assays

The intracellular growth assay of *L. pneumophila* was assayed as described previously (Wan et al., 2019). Briefly, *A. castellanii* was propagated using PYG medium. Cells were grown to near confluency and then plated into 24-well plates at a density of one million per well and were infected with stationary phase *L. pneumophila* at an MOI of 0.3 for 1 hr. The cells were then washed one time to remove extracellular bacteria and incubated at 37 °C for the indicated period. The amoeba cells were lysed with 0.05% saponin (Alfa Aesar, A18820) in PBS at 0, 20, 30, and 44 hrs, and the lysates were plated with serial dilutions onto CYE agar plates. Bacterial colonies were counted after 4 days of incubation at 37 °C. All growth assays were performed in triplicate.

## ACKNOWLEDGEMENTS

We thank Dr. Anthony Bretscher (Cornell Univ.) for providing beef skeletal muscle acetone powder for actin purification and plasmids (pRS415 yemCherry and pRS415 yeGFP); Dr. Scott Emr (Cornell Univ.) for yeast strain (SEY6210.1 with genomically tagged VPH1-mCherry). We also thank Dr. Joseph Vogel (Wash. Univ.) for the critical discussion. This work was supported by the National Institutes of Health (NIH) Grants R01-GM135379-01 (Y.M.). EM images were collected using an instrument supported by the NIH through award S10OD030470-01.

## AUTHOR CONTRIBUTIONS

Q.Z. and Y.M. conceived the project. Q.Z. and M.W. performed the experiments. Q.Z., M.W., and Y.M. analyzed the data. Q.Z., M.W., and Y.M. wrote the paper.

## SUPPLEMENTAL FIGURES and LEGENDS

**Figure 1—figure supplement 1.**
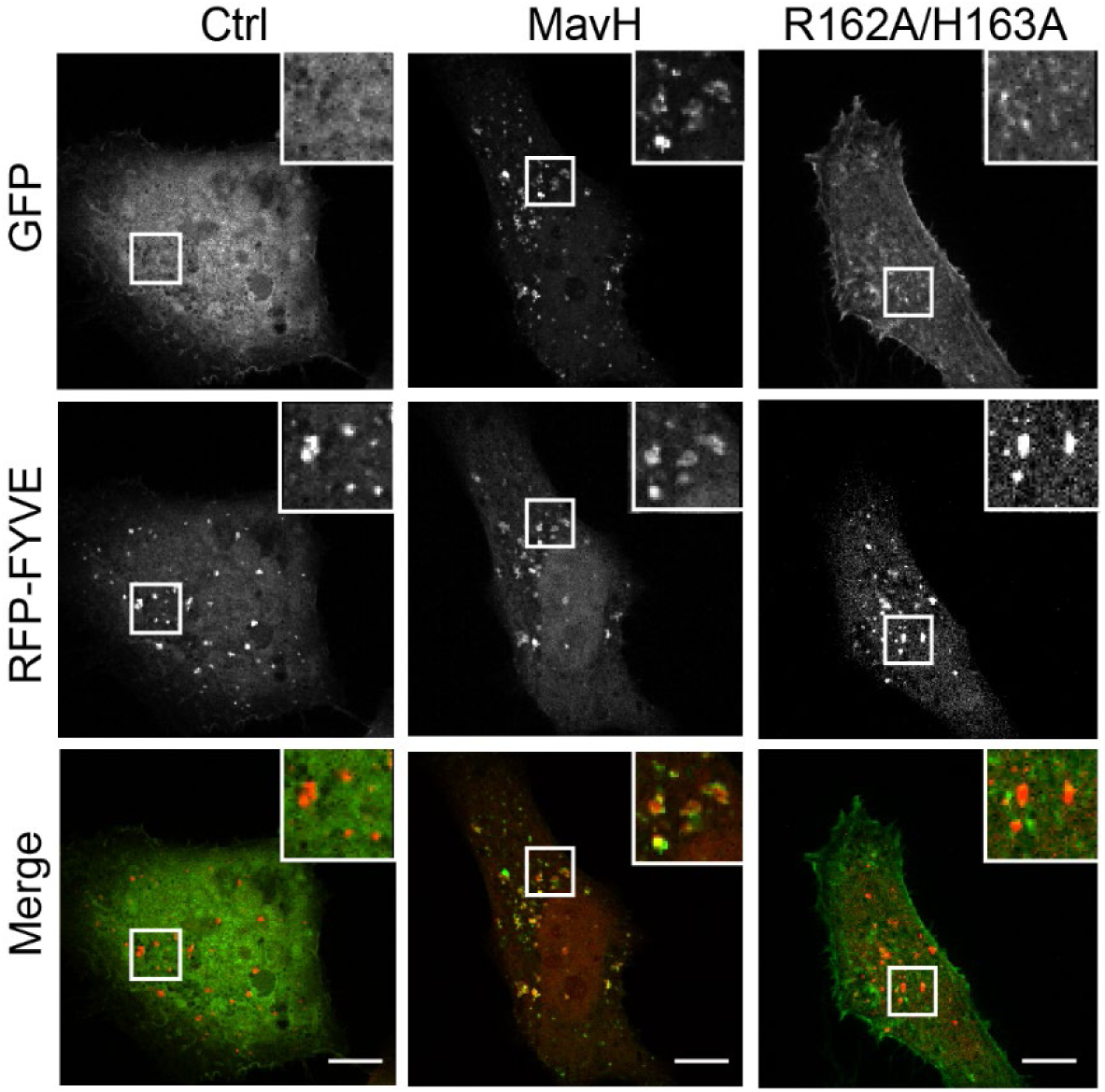
MavH co-localizes with the RFP-FYVE motif. Hela cells were co-transfected with a plasmid expressing RFP-2xFYVE with GFP, GFP-MavH, or GFP-MavH R162A/H163A for 20 hours. The colocalization of MavH with the PI(3)P marker RFP-2xFYVE was analyzed by fluorescence confocal microscopy. Wild-type MavH showed a high degree of colocalization with RFP-2xFYVE, while the MavH R162A/H163A mutant defective of PI(3)P-binding exhibited no colocalization with RFP-2xFYVE. Scale bars, 10 *µ*m.

**Figure 1—figure supplement 2.**
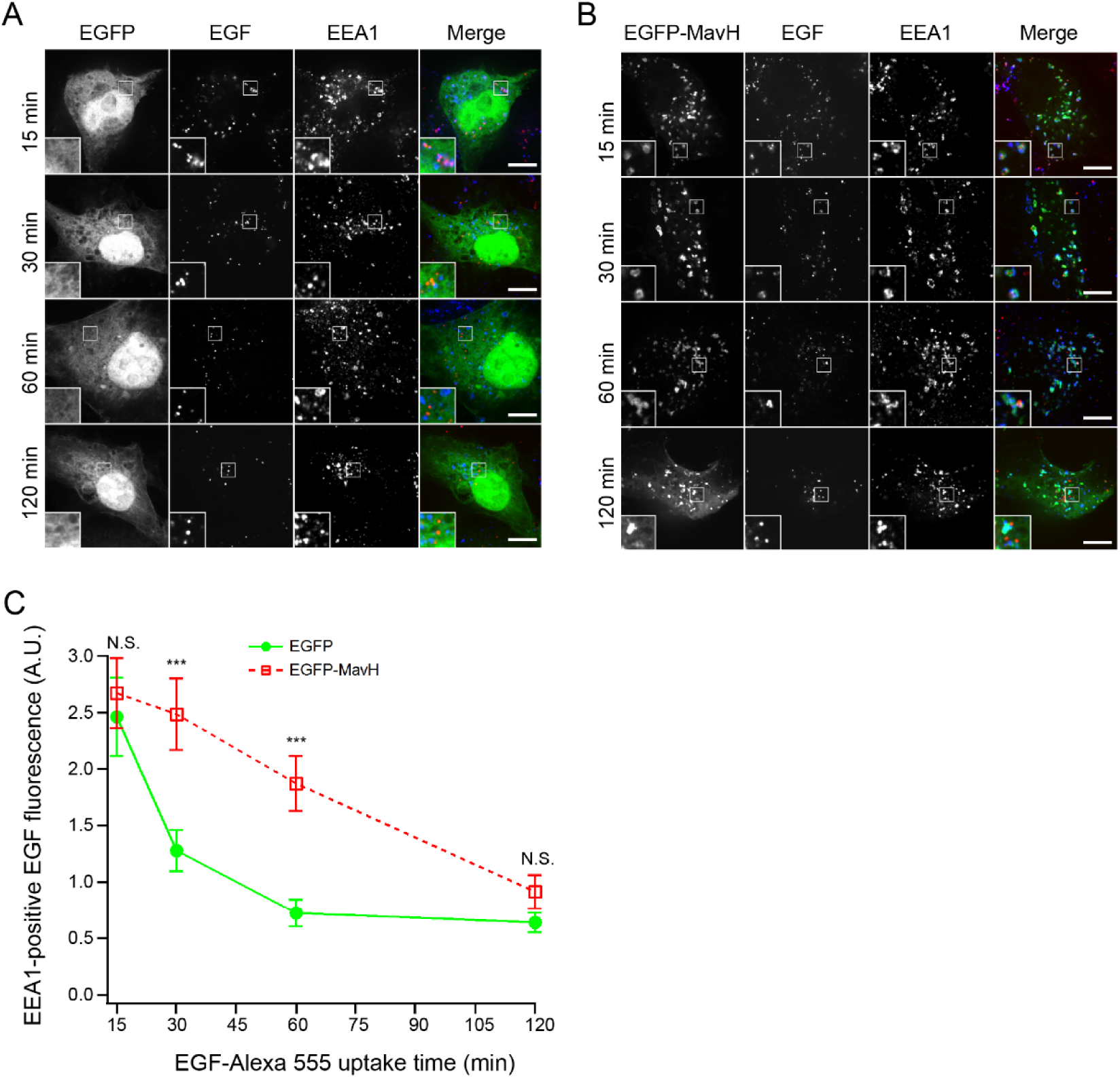
MavH inhibits endosomal trafficking. Cos7 cells were first transfected with indicated plasmids for 24 hours. Cells were then incubated with 20 ng/mL Alexa 555-EGF on ice for 20 min, washed, and then incubated at 37 °C for the indicated time. Early endosomes were stained with EEA1 antibodies. Representative images were shown for EGFP (A) or EGFP-MavH (B) transfected cells. EGFP-tagged protein was colored green, Alexa 555-EGF was colored in red, and EEA1 was colored in blue. Scale bars = 10 µm. (C) EEA1-positive EGF fluorescence was quantified, shown as Mean ± SEM from three independent experiments. At least 28 cells/condition were counted. ***P<0.001, N.S., not significant.

**Figure 1—figure supplement 3.**
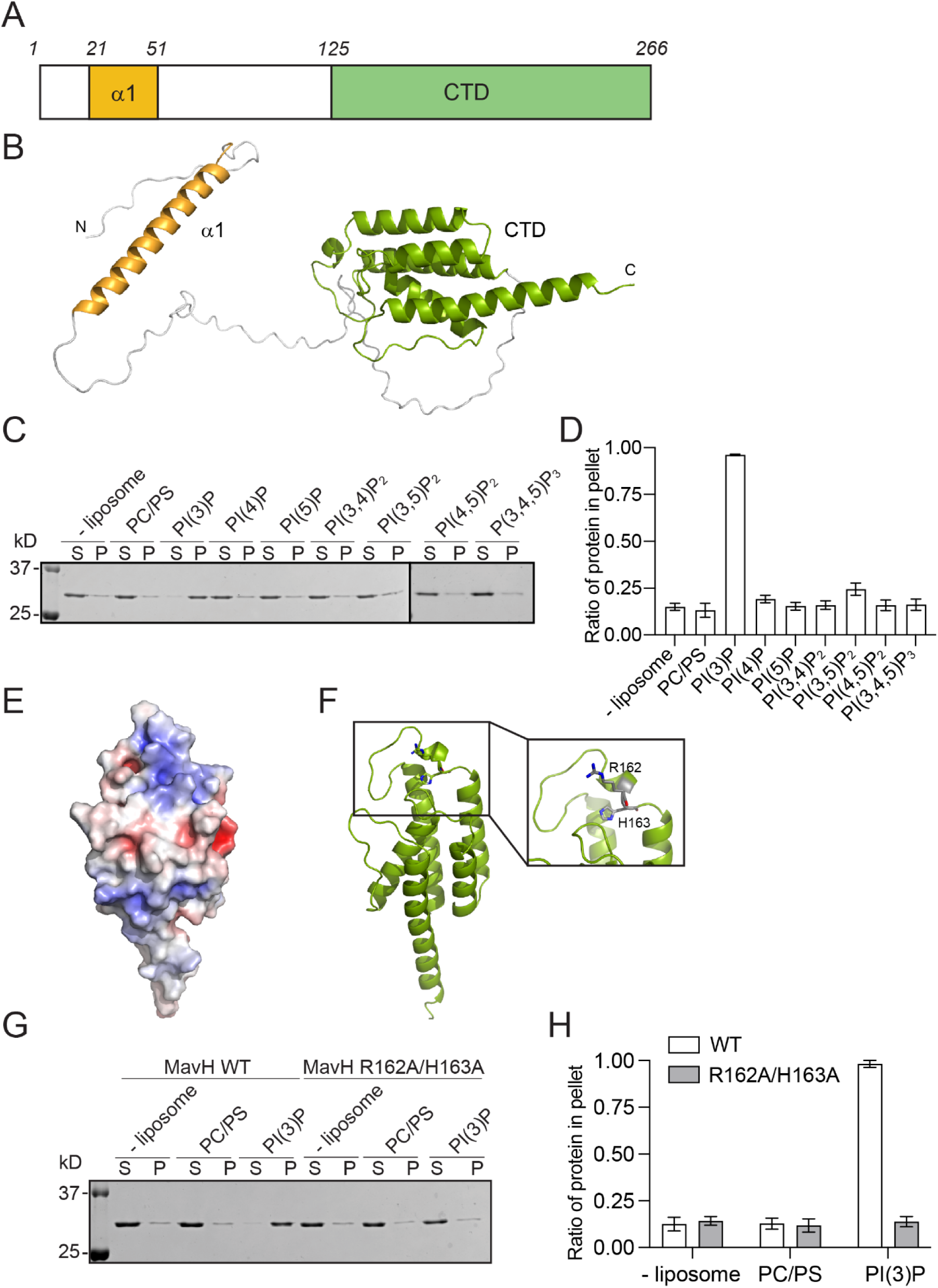
MavH binds PI(3)P via the C-terminal domain. (A) Schematic representation of MavH. N terminal *α* helix (*α*1) is colored in yellow. The C-terminal domain (CTD) is in green. (B) Ribbon diagram of predicated MavH structure with AlphaFold2. (C) Liposome co-sedimentation assays of MavH. Liposomes were formed with PC, PS, and indicated phosphoinositides. After incubation with MavH, the liposomes were pelleted by ultracentrifugation. P, pellet; S, supernatant. Pellet and supernatant fractions were then analyzed by SDS-PAGE, followed by Coomassie staining. (D) Quantification of the liposome sedimentation assays in (C). The ratios of the protein in the pellet were shown as Mean ± SEM from three independent experiments. (E) Molecular surface of CTD of MavH. The surface is colored based on electrostatic potential with the positively charged region in blue and the negatively charged surface in red. (F) Ribbon representation of CTD of MavH. The conserved positively charged residues, R162 and H163 are shown in sticks. (G) Liposome co-sedimentation assays of MavH wild type and R162A/H163A mutant. (H) Quantification of liposome sedimentation assays in (G). The ratios of the protein in the pellet were shown as Mean ± SEM from three independent experiments.

**Figure 3—figure supplement 1.**
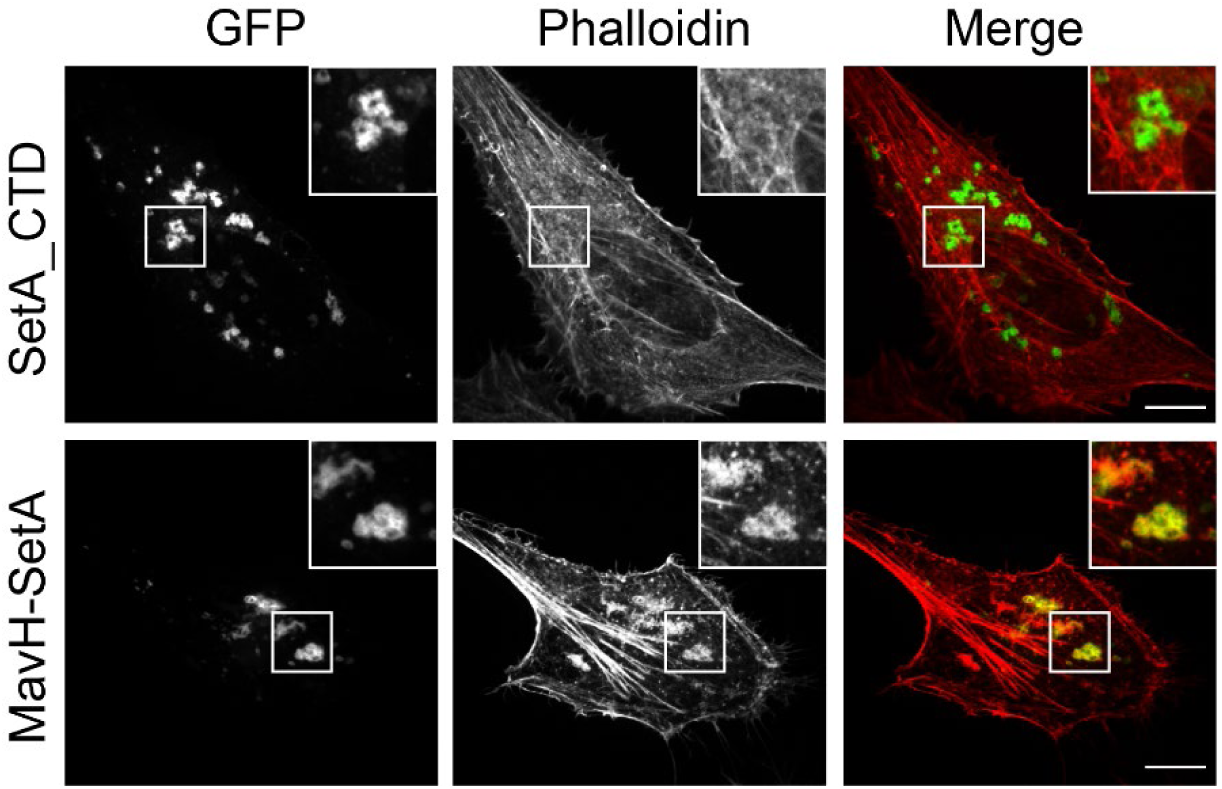
A chimeric fusion of MavH promotes actin polymerization around endosomes. Hela cells were transfected with GFP tagged SetA PI(3)P binding domain alone or a chimeric fusion containing the N-terminal portion of MavH in-frame fused with the SetA PI(3)P-binding domain. After transfection with indicated plasmids for 18 hrs, cells were fixed and stained with Rhodamine conjugated phalloidin and imaged by confcal microscopy. Scale bars = 10 µm.

**Figure 3—figure supplement 2.**
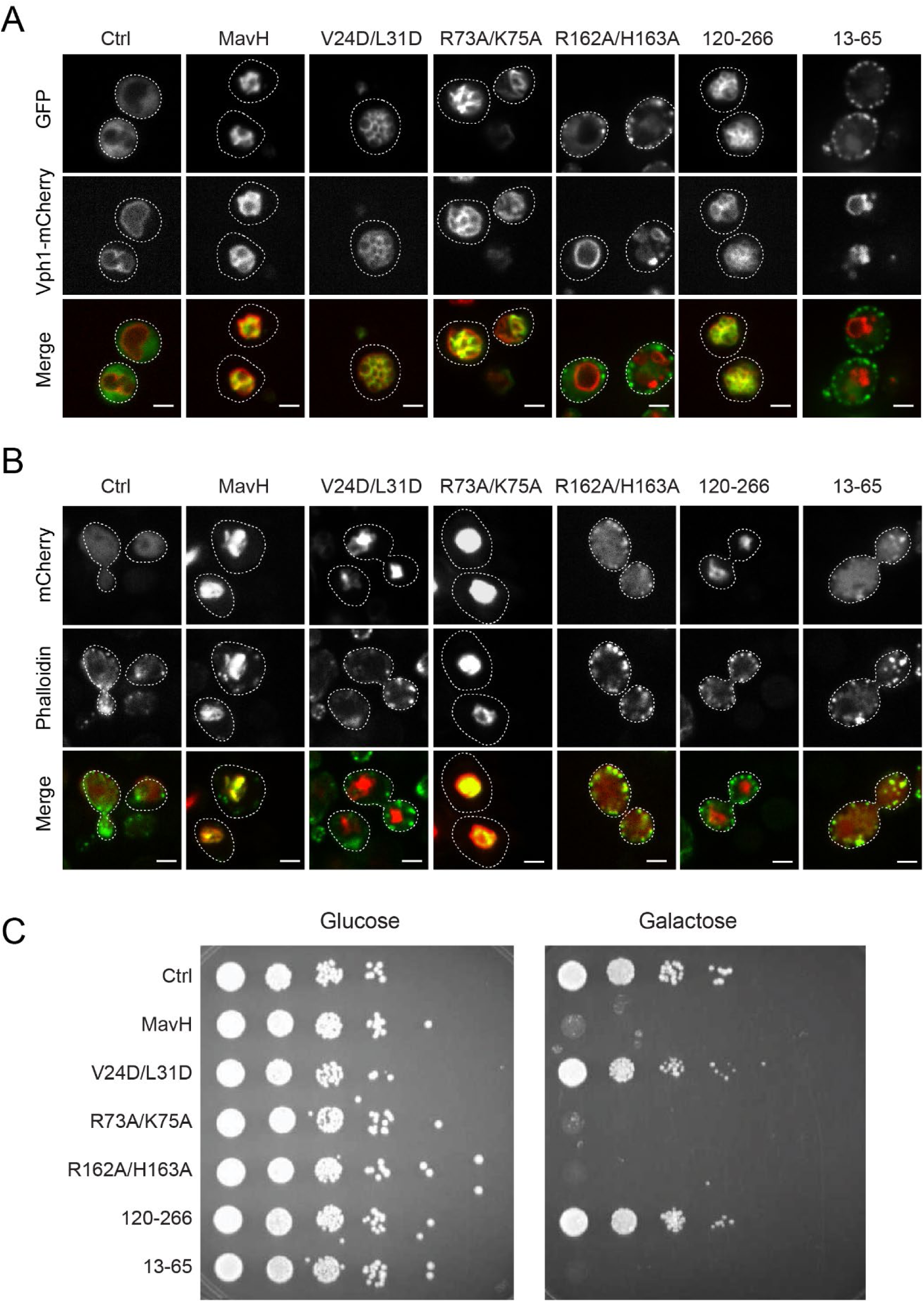
Intracellular localization and function of MavH in *S. cerevisiae.* (A) SEY6210.1 yeast strains expressing integrated copies of VPH1-mCherry were transformed with either GFP-tagged wild-type MavH, truncations, or mutants constructs under the control of a galactose inducible promoter. Cells were visualized by fluorescence confocal microscopy after induction of protein expression by selective media containing 2% galactose. (B) BY4741 yeast strains were transformed with either mCherry tagged wild-type or mutant MavH constructs under the control of a galactose inducible promoter. After induction of protein expression by selective media containing 2% galactose, yeast cells were stained with 488-phalloidin and visualized by fluorescence confocal microscopy. (C) Yeast cultures from (B) were grown on plates containing glucose or galactose (inducing conditions). 10 fold serial dilutions of each yeast cell culture were spotted on the plate and the lethal effects were compared to the empty vector control.

